# Reconstruction and analysis of thermodynamically-constrained models reveal metabolic responses of a deep-sea bacterium to temperature perturbations

**DOI:** 10.1101/2022.07.01.498526

**Authors:** Keith Dufault-Thompson, Chang Nie, Huahua Jian, Fengping Wang, Ying Zhang

## Abstract

Microbial acclimation to different temperature conditions can involve broad changes in cell composition and metabolic efficiency. A systems-level view of these metabolic responses in non-mesophilic organisms, however, is currently missing. In this study, thermodynamically-constrained genome-scale models were applied to simulate the metabolic responses of a deep-sea psychrophilic bacterium, *Shewanella psychrophila* WP2, under suboptimal (4°C), optimal (15°C), and supraoptimal (20°C) growth temperatures. The models were calibrated with experimentally determined growth rates of WP2. Gibbs free energy change of reactions (*Δ_r_G’*), metabolic fluxes, and metabolite concentrations were predicted using random simulations to characterize temperature-dependent changes in the metabolism. The modeling revealed the highest metabolic efficiency at the optimal temperature, and it suggested distinct patterns of ATP production and consumption that could lead to the lower metabolic efficiency under suboptimal or supraoptimal temperatures. The modeling also predicted rearrangement of fluxes through multiple metabolic pathways, including the glycolysis pathway, Entner-Doudoroff pathway, tricarboxylic acid (TCA) cycle, and the electron transport system, and these predictions were corroborated through comparisons to WP2 transcriptomes. Furthermore, predictions of metabolite concentrations revealed the potential conservation of reducing equivalents and ATP in the suboptimal temperature, consistent with experimental observations from other psychrophiles. Taken together, the WP2 models provided mechanistic insights into the metabolism of a psychrophile in response to different temperatures.

**Importance:** Metabolic flexibility is a central component of any organism’s ability to survive and adapt to changes in environmental conditions. This study represents the first application of thermodynamically-constrained genome-scale models in simulating the metabolic responses of a deep-sea psychrophilic bacterium to varying temperatures. The models predicted differences in metabolic efficiency that were attributed to changes in metabolic pathway utilization and metabolite concentration during growth under optimal and non-optimal temperatures. Experimental growth measurements were used for model calibration, and temperature-dependent transcriptomic changes corroborated the model-predicted rearrangement of metabolic fluxes. Overall, this study highlights the utility of modeling approaches in studying the temperature-driven metabolic responses of an extremophilic organism.

## Introduction

Temperature has a major impact on microbial physiology, affecting growth through physicochemical changes, such as altered enzyme stability (1–3), reaction kinetics (1), and membrane fluidity (4). Mesophilic organisms are known to respond to non-optimal growth temperatures using a variety of strategies such as the induction of cold or heat shock proteins (5–9), alteration of gene expression patterns (10), adjustment of membrane lipid composition (4), modification of protein synthesis and degradation (11, 12), and broad remodeling of metabolic pathways (13–15). A common cellular response to changes in temperature is the alternation of ATP levels in the cell, though variable responses have been seen among species adapted to different environments (16, 17). Additionally, the ability of bacteria to utilize ATP efficiently during the production of proteins and other biomass components has been shown to vary greatly in different conditions and at different growth rates (18, 19). Changes in temperature can also affect biogeochemical cycling by driving changes in the carbon use efficiency (CUE) of microbial communities (20). However, the influence of temperature on CUE is variable between different microbes and even within the same taxa (21). Overall, these prior studies demonstrate that responses to temperature changes can involve multiple physiological changes and be highly diverse among organisms.

Compared to the extensive understanding of temperature-related physiological responses in mesophiles, a comprehensive view of these processes is still being developed for psychrophiles, organisms having maximum growth temperatures around 20°C and optimal growth temperatures between 5°C and 15°C. Unlike the mesophiles, some psychrophiles constitutively express proteins commonly involved in cold shock responses (22, 23). A variety of metabolic responses can be induced in psychrophilic or psychrotolerant organisms under low-temperature conditions (24), including the differential expression of genes related to electron transport, carbon utilization, and energy production, along with the upregulation of the biosynthesis of branched-chain amino acids and unsaturated fatty acids (24–28). At elevated temperatures, psychrophiles have been observed to respond through the upregulation of heat shock proteins (29) and through changes in the regulation of energy metabolism (16, 30, 31).

The psychrophilic bacterium, *Shewanella psychrophila* WP2 (hereafter referred to as WP2), was isolated from benthic sediments in the western Pacific Ocean, where the ambient environmental temperature was approximately 4°C (32). WP2 has an optimal growth temperature of 10–15°C, a growth range of 0–20°C, and exhibits the hallmark metabolic versatility seen in the *Shewanella* genus (32, 33). WP2 belongs to the Group 1 *Shewanella*, a clade mainly consisting of deep-sea strains characterized by their tolerance and adaptation to low-temperature and high-pressure conditions (34, 35). WP2 represents a distinct clade that is distant from two other Group 1 *Shewanella* strains that have been represented using Genome-scale models (GEMs): *Shewanella loihica* PV-4 and *S. piezotolerans* WP3 (34, 36). An analysis of the complete genome of WP2 revealed an expansion of transposable elements, motility genes, and chemotaxis genes compared to other deep-sea *Shewanella* (37). However, so far little is known about the mechanisms of its psychrophilic adaptation or its physiological responses to non-optimal temperatures.

Genome-scale metabolic modeling has become a common technique for investigating genotype-phenotype associations in bacteria. Advancements in the development of GEMs have enabled their application in mapping metabolic responses of a growing number of extremophilic organisms under diverse environmental conditions (38–40). Constraint-based approaches are commonly used to simulate the utilization of metabolic pathways in GEMs, where optimal solutions of an objective function (*e.g.*, biomass production by a microorganism) are explored within the boundary of predefined metabolic constraints (*e.g.*, the stoichiometric constraints of biochemical reactions) (41, 42). Extensions of typical constraint-based approaches are often centered around the definition of additional constraints and exploration of the solution space. For example, constraints related to the thermostability of metabolic enzymes can be introduced based on the analyses of three-dimensional protein structures, leading to the identification of growth-limiting metabolic steps of a mesophilic bacterium under supraoptimal temperatures (43). Thermodynamics-based metabolic flux analysis (TMFA) uses thermodynamic constraints to inform the feasibility of reactions, permitting the incorporation of additional variables, including temperature, metabolite concentrations, and Gibbs free energy change of reactions (*Δ_r_G’*), into the model representation (44, 45). While changes in thermodynamic favorability due to temperature are known to have effects on metabolic pathway utilization (46–48), methods like TMFA have not been extensively applied to simulate metabolism at different temperatures.

To build towards a systems-level understanding of the metabolic responses of psychrophilic bacteria to different temperatures, thermodynamic constraints were applied during the modeling of temperature-dependent metabolic changes in WP2, a deep-sea psychrophilic bacterium. The predicted metabolic pathway utilization and metabolic efficiency measures were compared among different growth temperatures to investigate the underlying mechanisms involved in temperature acclimation in WP2.

## Results

### Temperature-dependent, thermodynamically constrained genome-scale models of WP2

A genome-scale metabolic reconstruction was developed following the iterative annotation of the WP2 genome (NCBI accession: GCA_002005305.1) (**Materials and Methods**). The resulting WP2 GEM had an overall consistency score of 99% based on the MEMOTE test suite (49) and contained 940 reactions, 786 genes, and 683 metabolites (**Data Set S1**). It represents experimentally confirmed growth and no-growth phenotypes on combinations of carbon sources (*e.g.*, glucose, galactose, cellobiose, etc.) and electron acceptors (*e.g.*, nitrate, trimethylamine N-oxide, Fe(III), etc.) (**Data Set S1, Tab 1**). The construction of temperature-dependent, thermodynamically constrained models was performed using the TMFA approach (**Materials and Methods**). Thermodynamic constraints were included for 89% (839 reactions) of the 940 reactions included in the WP2 GEM, with the rest of the reactions either representing macromolecular biosynthesis or involving complex metabolites that have unknown Gibbs free energies of formation (**Data Set S1**). Temperature dependency was introduced through the calculation of Gibbs free energy change of reactions (*Δ_r_G’*) for the representation of models under optimal (15°C), supraoptimal (20°C), and suboptimal (4°C) temperatures (**Materials and Methods**).

Each temperature-dependent model was calibrated by using experimentally measured growth rates to constrain an ATP hydrolysis reaction (ATPM) that accounts for the non-growth associated ATP utilization by a cell. Briefly, this ATP maintenance flux was determined by fitting the model predictions to experimentally measured growth rates (**Fig. 1A**), and the constraint range of the ATP maintenance flux was determined based on the experimentally derived growth rates of three biological replicate cultures of WP2 grown at each temperature (**Materials and Methods**). The lowest ATP maintenance cost was observed under 15°C, with a calibrated ATP hydrolysis constraint ranging between 1.32 and 1.36 (**Fig. 1B**). Compared to the 15°C condition, the calibrated ATP maintenance constraint was doubled under 4°C, ranging between 2.71 and 2.82 (**Fig. 1C**). The 20°C model had much higher ATP maintenance cost and also higher variability in the calibrated ATP hydrolysis constraint compared to the 15°C or 4°C models, ranging between 7.41 and 11.55 (**Fig. 1D**). This aligned with the lower growth rate and a greater level of growth variability (as measured by the standard deviations of the growth yields under each sampling point) across experimental replicates at 20°C. The ATP maintenance constraints were applied for all subsequent simulations of the corresponding temperature-dependent models unless otherwise noted.

**Fig. 1.**
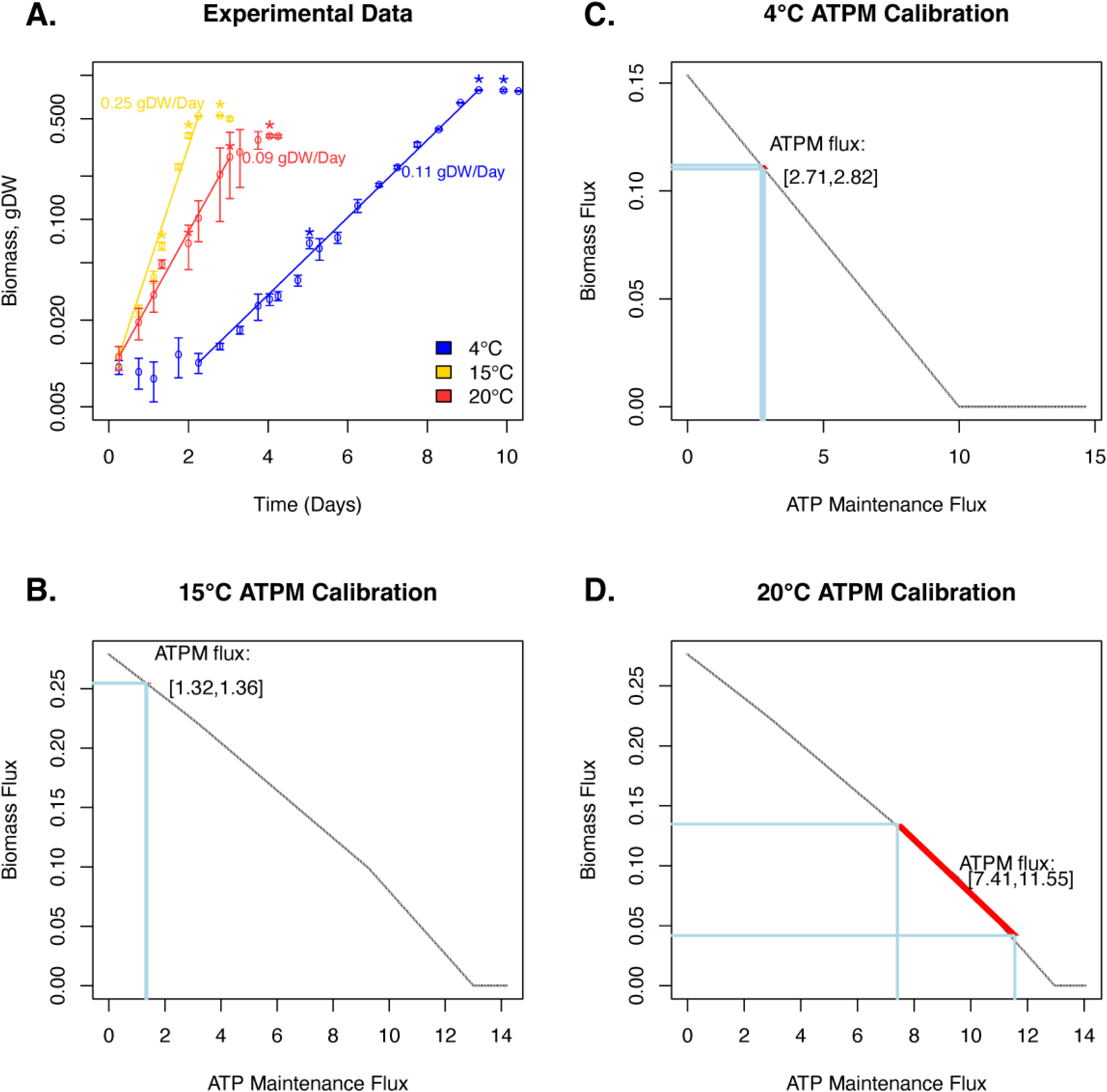
Temperature-dependent calibration of metabolic constraints on the non-growth associated ATP maintenance. **A.** Experimental measurements of WP2 growth under 4°C (blue), 15°C (yellow), and 20°C (red). The open circles and error bars represent the average and standard deviation of growth yields, measured at each time point over three biological replicates. The solid lines indicate periods of exponential growth at each condition. Labels indicate the average growth rate from each temperature over three biological replicates. Asterisks indicate timepoints where RNA-Seq samples were taken. **B-D.** Calibration of the ATPM fluxes in the 15°C **(B)**, 4°C **(C)**, and 20°C **(D)** models based on experimental data. Successively increasing ATPM fluxes (x-axis) were plotted against the optimized biomass fluxes (y-axis) determined from the robustness simulation. Horizontal lines were used to indicate the range of experimentally determined growth rates. Vertical lines were used to indicate the range of calibrated ATPM fluxes.

### Temperature-dependent changes in metabolic efficiency

The temperature-dependent models were simulated with N-acetyl D-glucosamine (GlcNac) as a sole carbon source in aerobic conditions, replicating the conditions used in the WP2 growth experiments (**Materials and Methods**). For each temperature-dependent model, 1,000 random simulations were performed with each simulation constrained by fixing the biomass flux to a random value selected within the range of experimentally derived growth rates (**Materials and Methods**). Predictions of reaction fluxes, Gibbs free energy change of reactions (*Δ_r_G’*), and metabolite concentrations were generated from each random simulation and were used to characterize the metabolic changes between different temperatures.

The metabolic efficiency was estimated based on each random simulation using three global efficiency parameters: CUE, ATP production per carbon substrate (ATP produced / GlcNac), and ATP consumption per gram dry weight (gDW) of biomass (**Materials and Methods**). Of the three temperature-dependent models, the 15°C model showed the highest metabolic efficiency, with a high CUE (2- to 3-fold higher than what was predicted in 4°C or 20°C models), a high ATP production, and a low ATP consumption in biomass synthesis (**Fig. 2**). While the 4°C and 20°C models had similar CUE, they showed distinct profiles in both ATP production and ATP consumption. The 20°C model was characterized by a high level of metabolic variability. It had an overall higher ATP production and higher ATP consumption compared to the other two models. The 4°C model, in contrast, demonstrated the lowest ATP production per carbon substrate. The predicted ATP consumption in biomass synthesis under 4°C was higher than the 15°C model but on average lower than the 20°C model (**Fig. 2**).

**Fig. 2.**
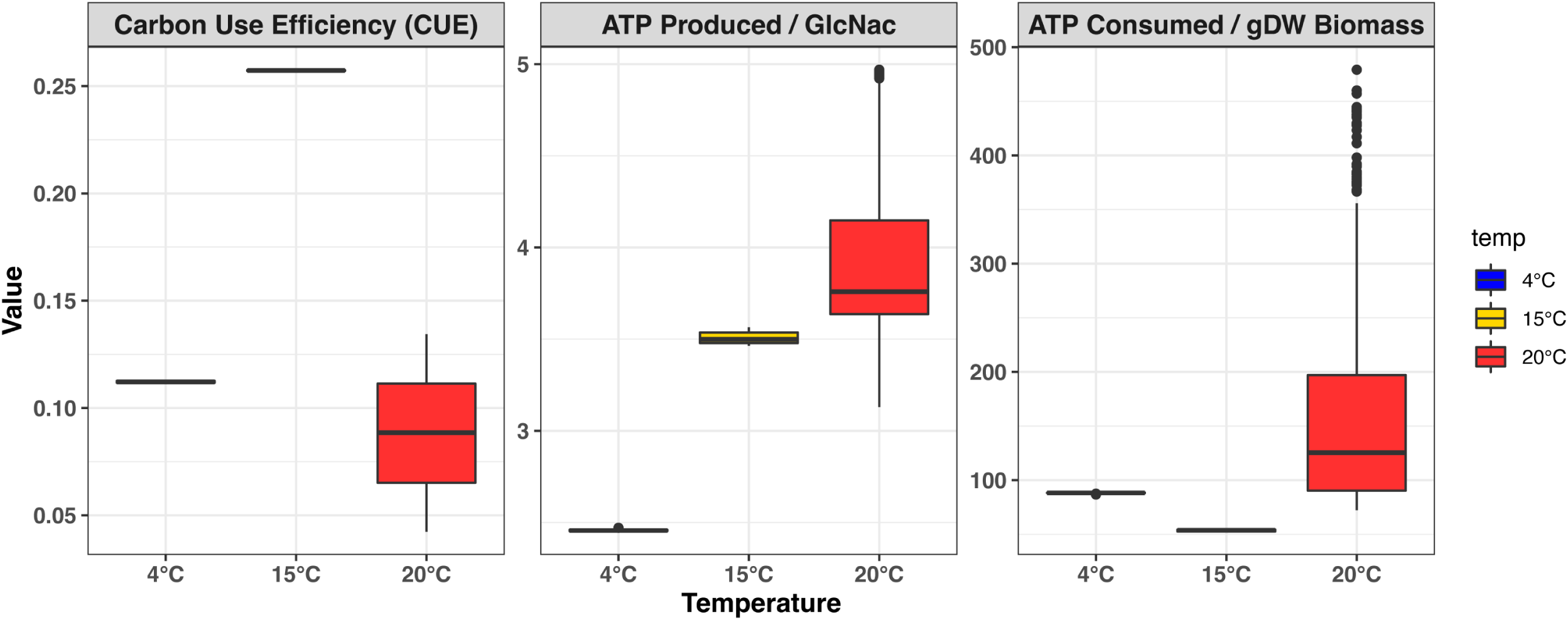
Predictions of metabolic efficiency by the temperature-dependent models. Boxplots showing the carbon use efficiency **(A)**, ATP produced per unit of carbon source **(B)**, and ATP consumed per gram dry weight of biomass **(C)** predicted by the 4°C (blue), 15°C (yellow), and 20°C (red) models based on 1,000 random simulations.

### Temperature-dependent remodeling of metabolic pathway usage

Comparisons of the Gibbs free energy change of reactions (*Δ_r_G’*) and reaction fluxes predicted from the 1,000 random simulations revealed a broad, temperature-dependent remodeling of metabolic pathway utilization in WP2. Of the 940 metabolic reactions, significant changes in metabolic flux were identified for 326 (35%) reactions and significant changes in *Δ_r_G’* were observed in 259 (28%) reactions (**Data Set S2**).

Multiple differences were observed in the utilization of central metabolic pathways under the different temperatures (**Fig. 3**). One of the main differences was related to how the temperature-dependent models utilize the payoff phase of glycolysis, which catalyzes the chemical conversion from glyceraldehyde 3-phosphate (G3P) to phosphoenolpyruvate (PEP), generating cellular energy in the form of ATP and reduced forms of electron carriers (i.e., NADH). The pathway was predicted to carry flux in both the 15°C and 20°C models; however, it was blocked under 4°C due to the glyceraldehyde-3-phosphate dehydrogenase (GAPD) reaction being thermodynamically unfavorable at the lower temperature. Instead, the carbon flow in the 4°C model was redirected through the gluconeogenic reactions, fructose bisphosphate aldolase (FBA), and fructose bisphosphatase (FBP), towards the oxidative pentose phosphate pathway (PPP) and the Entner-Doudoroff (ED) pathway. The utilization of the oxidative PPP and ED pathways was optional in the 15°C or 20°C models.

**Fig. 3.**
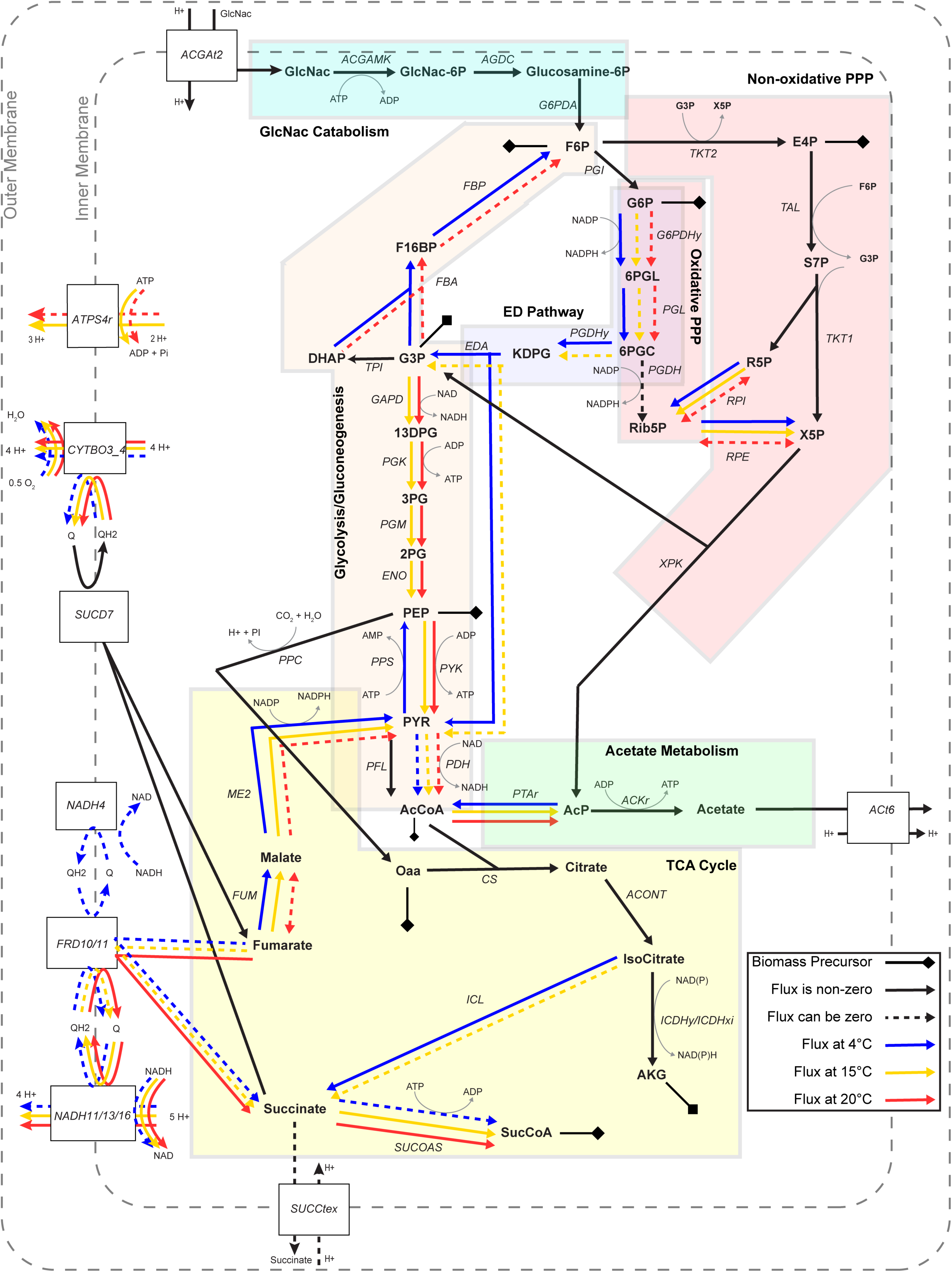
Temperature-dependent utilization of metabolic pathways. Pathway diagram showing the direction of metabolic fluxes predicted by the different models. Solid lines indicate reactions that are obligated to carry non-zero fluxes in the shown direction, while dashed lines indicate that the reaction flux was zero in some instances of the 1,000 random simulations. Black lines represent common pathways used by WP2 under all three temperatures. Colored lines represent pathways used under 4°C (blue), 15°C (yellow), or 20°C (red). Metabolic pathways were shown with background shades and labeled by the pathway name. Precursor metabolites for biomass production are marked with a line capped with a diamond. Abbreviations: GlcNac: N-acetyl-D-glucosamine, GlcNac-6P: N-acetyl-D-glucosamine 6-phosphate, Glucosamine-6P: D-glucosamine 6-phosphate, F6P: D-fructose 6-phosphate, F16BP: D-fructose 1,6-bisphosphate, DHAP: dihydroxyacetone phosphate, G3P: glyceraldehyde 3-phosphate, 13DPG: 3-phospho-D-glyceroyl phosphate, 3PG: 3-phospho-D-glycerate, 2PG: D-glycerate 2-phosphate, PEP: phosphoenolpyruvate, PYR: pyruvate, AcCoA: acetyl-CoA, G6P: D-glucose 6-phosphate, 6PGL: 6-phospho-D-glucono-1,5-lactone, 6PGC: 6-phospho-D-gluconate, Rib5P: D-ribulose 5-phosphate, KDPG: 2-dehydro-3-deoxy-D-gluconate, E4P: D-erythrose 4-phosphate, S7P: sedoheptulose 7-phosphate, R5P: alpha-D-ribose 5-phosphate, X5P: D-xylulose 5-phosphate, AcP: acetyl phosphate, Acetate: acetate, Oaa: oxaloacetate, Citrate: citrate, IsoCitrate: isocitrate, AKG: 2-oxoglutarate, SucCoA: succinyl-CoA, Succinate: succinate, Fumarate: fumarate, Malate: L-malate, ATP: adenosine triphosphate, ADP: adenosine diphosphate, Pi: orthophosphate, H+: proton, NADH: nicotinamide adenine dinucleotide (reduced), NAD: nicotinaminde adenine dinucleotide, NADPH: nicotinamide adenine dinucleotide phosphate (reduced), NADP: nicotinamide adenine dinucleotide phosphate, QH2: quinol pool, Q: qinone pool

The 4°C model also differed from the 15°C or 20°C models in its utilization of the phosphate acetyltransferase (PTAr) reaction (**Fig. 3**). PTAr was thermodynamically constrained to carry flux in the acetyl-phosphate (AcP) to acetyl-CoA (AcCoA) direction in the 4°C model, likely contributing to carbon conservation through the production of AcCoA. In contrast, the PTAr reaction in the 15°C and 20°C models carried flux in the AcCoA-consuming direction and was linked to the downstream production of ATP and acetate via the acetate kinase reaction (ACKr).

The three temperature-dependent models also varied in their utilization of the tricarboxylic acid (TCA) cycle. The 4°C model used the isocitrate lyase (ICL), fumarase (FUM), and malic enzyme (ME2) reactions, conserving carbon for the production of pyruvate (PYR), which in turn could be used for the synthesis of the precursor metabolites, PEP and AcCoA (**Fig. 3**). The FUM and ME2 reactions were used in the 15°C model but were optional in the 20°C model, while the ICL reaction was not used under 20°C and was optional under 15°C.

Outside of the central carbon metabolism, differences were also observed in the modeled utilization of the electron transport system. The ATP synthase reaction (ATPS4r) was used in the proton-pumping direction in the 15°C model and was optional in the same direction in the 20°C model. However, ATPS4r was not used in the 4°C model. The 20°C model obligately used the proton-translocating NADH:quinone oxidoreductase reactions (NADH 11/13/16) to carry out electron transport from NADH to the quinone pool. This was connected to the fumarate reductase reaction (FRD 10/11), apparently contributing to the cycling of electron carriers in the 20°C model. In contrast, the 4°C model could utilize the alternative, non-proton-translocating NADH dehydrogenase reaction (NADH4), which was not used in either the 15°C or the 20°C model (**Fig. 3**). Overall, higher variability and lower fluxes were seen in the electron transport reactions (e.g., SUCD7, CYTBO3_4, FRD10/11, and NADH11/13/16) in the 4°C model compared to the 15°C or 20°C models ( **Fig. S1**).

The remodeling of electron transport pathways was reflected in the changes of the overall redox potential, measured by the ratios between concentrations of reduced versus oxidized forms of electron carrier metabolites (**Fig. 4**). The 4°C model demonstrated the highest [NADH]/[NAD^+^] and [NADPH]/[NADP^+^] ratios, which may indicate the conservation of reducing equivalents and was consistent with the reduced metabolic fluxes in the electron transport system, such as the NADH:quinone oxidoreductase (**Fig. S1**). Surprisingly, despite a predicted low ATP production efficiency and a high ATP consumption for biomass synthesis in the 4°C model (**Fig. 2**), a higher [ATP]/[ADP] ratio was observed in the 4°C model compared to the 15°C or the 20°C models (**Fig. 4**), suggesting a high level of energy conservation at the low temperature.

**Fig. 4.**
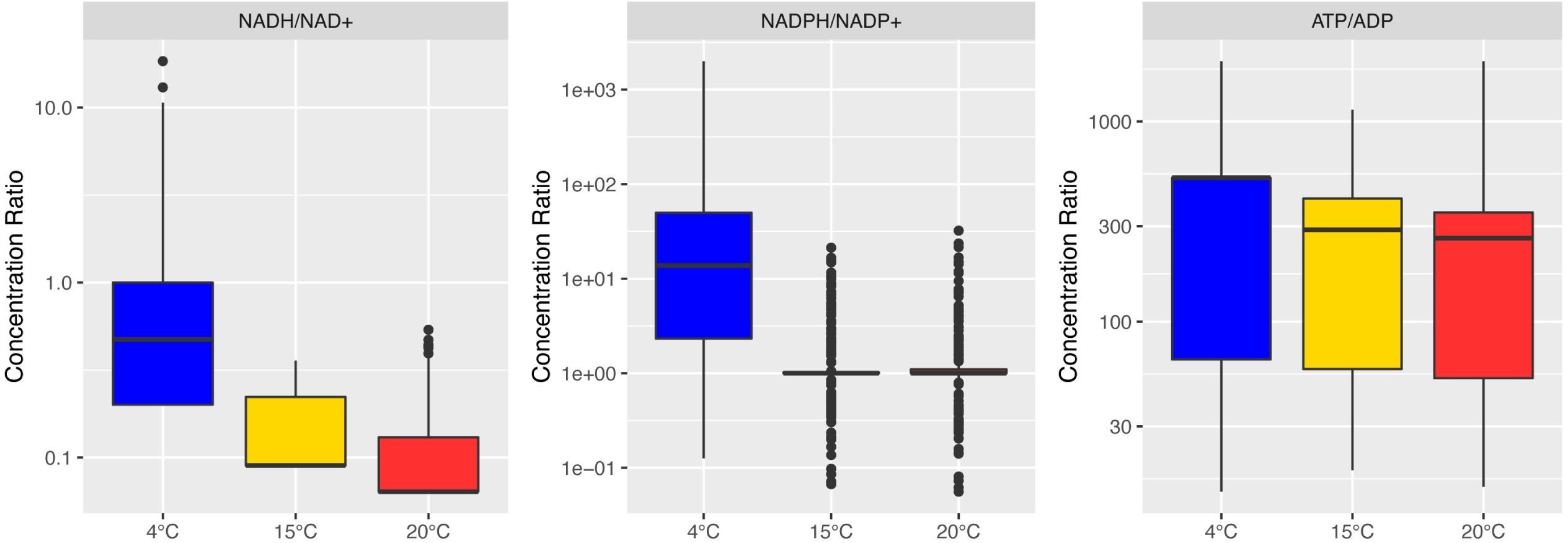
Ratios of metabolite concentrations predicted by the temperature-dependent models. Ratios of [NADH]/[NAD^+^] **(A)**, [NADPH]/[NAPD^+^] **(B)**, and [ATP]/[ADP] **(C)** are shown as boxplots based on 1,000 random simulations for the 4°C (blue), 15°C (yellow), and 20°C (red) models.

### Differential expression of metabolic genes

The temperature-dependent metabolic responses of WP2 were experimentally characterized using transcriptomes obtained from three growth phases: early exponential, late exponential, and stationary (**Materials and Methods**). Comparisons of the transcriptomes revealed a very weak negative correlation between the expression of genes at 4°C compared to the other two temperatures, similar to what was observed on the model-predicted metabolic fluxes and *Δ_r_G’* values (**Fig. S2**). Analyses of the expression of genes associated with specific metabolic reactions showed a great level of variability across different growth phases (**Data Set S3**) but also revealed agreement between the differential expression of genes and the temperature-dependent differences in fluxes of some key reactions from the carbon metabolism and electron transport pathways, especially during the late exponential growth phase (**Fig. S1**).

The pyruvate formate lyase (PFL) gene was significantly upregulated under 15°C in the late exponential phase. This was consistent with the high metabolic flux predicted for the PFL reaction and the prediction of a more active glycolytic pathway, an upstream pathway to PFL, in the 15°C model. The expression of the PFL gene at 20°C was higher than 4°C during the early exponential phase but lower than 4°C during the late exponential or stationary phases. This was in line with the high variability of PFL metabolic fluxes in the 20°C model.

Other examples of differentially expressed genes were observed in the central carbon metabolism. Genes encoding the FBA, glucose 6-phosphate dehydrogenase (G6PDHy), and phosphogluconate dehydratase (PGDHY) reactions were upregulated under 4°C (**Fig. S1**). This was consistent with the model-predicted utilization of the oxidative PPP and ED pathways under 4°C (**Fig. 3**). The ICL gene was similarly upregulated under 4°C in the late exponential phase. This was also in line with the model-predicted use of ICL in the 4°C model. The PTAr gene was upregulated under 15°C and 20°C compared to 4°C, corresponding with the predicted higher magnitude of PTAr fluxes under these optimal and supraoptimal temperatures (**Fig. S1**).

Additional examples involved the differential expression of genes in the electron transport systems. Specifically, multiple genes encoding subunits of the ATP synthase and the proton-translocating NADH:quinone oxidoreductase were upregulated under 20°C compared to 4°C, and the non-proton-translocating NADH dehydrogenase was upregulated in 4°C compared to 20°C . These were consistent with the model predictions, as the 20°C model predicted higher fluxes through the NADH11/13/16 reactions, and the 4°C model predicted the potential usage of NADH4 and lack of utilization of ATPS4r (**Fig. S1**).

## Discussion

Genome-scale metabolic modeling has been applied to a variety of different biological problems, but the incorporation of complex environmental factors like pH, pressure, or temperature into simulations of growth remains challenging. In this study, thermodynamically constrained models were developed of WP2, a deep-sea psychrophilic bacterium, setting an example for modeling metabolic responses to different temperatures.

The temperature-dependent, thermodynamically constrained models were calibrated based on experimentally measured growth of WP2 under optimal (15°C), suboptimal (4°C) and supraoptimal (20°C) temperatures, suggesting the lowest ATP maintenance cost in the 15°C model and a 2- and 7-fold higher ATP maintenance cost in the 4°C and 20°C models, respectively (**Fig. 1**). Higher variability was predicted in the calibrated range of ATP maintenance fluxes in the 20°C model compared to the 15°C or 4°C models, which can be attributed to the greater variability among replicates of the growth experiments. Interestingly, the experimental replicates from 20°C revealed an earlier start of the exponential growth phase but a lower average growth rate compared to the suboptimal temperature of 4°C (**Fig. 1**). This may indicate potential growth deficiency under 20°C, likely due to high-temperature stress, as 20°C represents the highest temperature that WP2 is known to grow at (33).

To provide a quantitative estimation of the metabolic efficiency, three global parameters, including the CUE, the ATP production per carbon substrate, and the ATP consumption per gDW of biomass synthesis, were calculated for each temperature-dependent model based on metabolic fluxes from 1,000 rounds of random simulations (**Materials and Methods**). These global parameters revealed the highest metabolic efficiency in the 15°C model, characterized by high CUE, high ATP production per carbon substrate, and low ATP consumption compared to the other two temperatures. The predicted CUE under 15°C had a median value of 0.26 (**Fig. 2**). This was similar to values predicted for other bacteria in substrate-limited growth conditions (50), indicating similar levels of metabolic efficiency in WP2 during optimal growth.

The two models of non-optimal temperatures (4°C and 20°C) predicted similar CUE but distinct measures of ATP production and ATP consumption (**Fig. 2**). The 4°C model had low efficiency in ATP production and moderately high ATP consumption in biomass synthesis, while the 20°C model had high ATP production and even higher ATP consumption in biomass synthesis compared to the 4°C model. It is worth mentioning that our calculation of the ATP consumption did not include the non-growth associated ATP maintenance (ATPM) for each temperature-dependent model. Therefore, the predicted high ATP consumptions in 4°C and 20°C models should reflect the high metabolic demands for each unit of biomass synthesis under these non-optimal temperatures.

The suboptimal temperature modeled in this study (4°C) represents the ambient temperature of WP2 in the deep sea (33). Our modeling result may indicate that WP2 is characterized by slow growth, with low CUE and low ATP production in its *natural* environment. This is similar to the slow growth phenotypes that have been observed in other deep-sea microbes (51, 52). Interestingly, slow growth has also been proposed as a survival strategy in bacteria, allowing them to better survive in oligotrophic or other extreme conditions (53–55). The 20°C model, while representing the highest tolerable temperature for WP2, presented the lowest growth rate and highest variability in the model-predicted metabolic fluxes. It has been suggested that this kind of flux variability may be related to a growth-flexibility trade-off, where slower growth may facilitate the use of alternative metabolic pathways while adjusting to environmental changes (56).

The temperature-dependent changes in metabolic efficiency may be linked to altered metabolic pathway usage predicted under the different temperatures. The higher ATP production in 15°C and 20°C models could be attributed to the utilization of the payoff phase of glycolysis (**Fig. 3**), as the ATP generating reactions in this pathway (e.g., PGK and PYK) were utilized in 15°C and 20°C models but blocked under 4°C due to unfavorable thermodynamics of the GAPD reaction (**Data Set S2**). The downstream reaction, PFL, accordingly carried higher fluxes in the 15°C and 20°C models compared to the 4°C model, consistent with the experimentally observed upregulation of the PFL gene under the higher temperatures (**Fig. S1**). The decrease of glycolytic fluxes in WP2 under 4°C was similar to the low-temperature responses observed in other psychrophilic bacteria (26, 27, 57), suggesting that the differential use of glycolysis may be driven by the thermodynamics of this pathway in low-temperature conditions.

The absence of glycolysis flux under 4°C led to the redirection of metabolic fluxes through the oxidative PPP and ED pathways. This was again consistent with the upregulation of genes associated with these pathways (e.g., FBA, G6PDHy, and PGDHY) (**Fig. S1**). It has been documented that the glycolytic pathways have tighter thermodynamic bottlenecks than the ED pathway (58), and the ED pathway is known to have a lower protein synthesis cost (59, 60). A similar metabolic strategy has also been reported in another psychrophilic gammaproteobacterium, *Colwellia psychrerythraea* (61)

Additional temperature-dependent alternations of metabolic pathway usage were observed in the TCA cycle and the electron transport chain (**Fig. 3**). The use of ICL reaction in the 4°C model was consistent with the upregulation of the ICL gene under 4°C in the late exponential phase (**Fig. S1**). Similarly, the upregulation of non-proton-translocating NADH dehydrogenase under 4°C aligned with the utilization of NADH4 reaction in the 4°C model (**Fig. S1**). The ICL reaction is considered to be a route for carbon conservation as it bypasses the CO_2_-producing steps in the TCA cycle (62). The upregulation of ICL has been observed under other stress conditions, such as oligotrophy or Fe-limitation, and has been linked to the reduction of electron transport activity (63, 64). This potentially aligns with the prediction of relatively lower metabolic fluxes carried by the NADH:quinone oxidoreductase (NADH11/13/16) in the 4°C model and is consistent with the gene expression data, especially in contrast to the 20°C condition (**Fig. S1**). Considering that our models predicted higher [NADH]/[NAD^+^] and [NADPH]/[NADP^+^] ratios under 4°C (**Fig. 4**), metabolic remodeling in the TCA cycle and electron transport systems may reflect potential shifts in the intracellular redox balance under the suboptimal temperature.

The ATPS4r was predicted to carry flux in the 15°C or the 20°C models, but not in the 4°C model (**Fig. 3**). Consistent with these temperature-dependent predictions, significant upregulation of genes encoding multiple subunits of the ATP synthase was observed in the late exponential phase when comparing transcriptomes in 15°C or 20°C to 4°C (**Data Set S3**). The ATPS4r reaction, when carrying non-zero fluxes, was predicted to run only in the ATP-consuming, proton-pumping direction (**Fig. 3**), similar to what has been observed in other *Shewanella* species (34, 65). The lack of ATPS4r flux in the 4°C model likely reflects a mechanism of ATP conservation. This was consistent with the higher [ATP]/[ADP] ratio predicted in the 4°C model compared to the 15°C or 20°C models (**Fig. 4**). It has been documented that psychrophilic organisms increase intracellular ATP levels in response to lower temperatures despite their reduced growth rates (66, 67). Our modeling results were consistent with these observations and suggested that WP2 may carry out similar responses to other psychrophiles under low temperatures.

Overall, our reconstruction of thermodynamically constrained genome-scale models provided an integrated view of the metabolic responses to different temperature conditions by a deep-sea psychrophilic bacterium. Several temperature-dependent changes in metabolic efficiency, pathway usage, and metabolite ratios were predicted by the models, demonstrating the complexity of metabolic responses under optimal and non-optimal temperatures. Future developments of this approach may include the incorporation of physiological changes associated with growth in different temperatures, such as altered membrane structures and changes in biomass composition. The modeling approach can be broadly applied to study temperature-dependent adaptations of other organisms, contributing to a systems-level understanding of metabolic responses to changing temperatures.

## Materials and methods

### WP2 growth experiments

Inoculations of three biological replicates for each growth temperature (4°C, 15°C, and 20°C) were performed in 150 mL of fresh LMO-812 media (34) with 1 mL of inoculum obtained from cultures grown to exponential phase at the optimal temperature (15°C) in 2216 Marine Medium (Difco). The cultures were grown with 5 mM GlcNac as the sole carbon source in aerobic conditions at each temperature with shaking, and an initial OD_600_ of 0.002 was measured for each inoculation. Turbidity was monitored throughout the growth phases of the cultures, and the OD_600_ values were converted to gram dry weight values for each culture using a previously established relationship between OD and dry weight biomass in *Shewanella* (*68*). Growth rates were calculated based on the periods of exponential growth for each temperature condition (**Fig. 1**).

### Transcriptome sampling and differential gene expression analysis

Samples for transcriptome sequencing were taken from the early exponential, late exponential, and stationary phases for each replicate in the 4°C, 15°C, and 20°C cultures (**Fig. 1**). Two biological replicates were sampled from each of the targeted growth phases for each temperature, where 2 mL of culture was taken from each replicate, centrifuged at 12000 times gravity for 2 minutes, and the cell pellet was used for the application of RNA extraction and sequencing using services provided by the Sangon Biotech in Shanghai, China. Total RNA was extracted from the cell pellets and cleaned using the Ribo-off rRNA depletion kit. cDNA libraries were prepared using the VAHTS Stranded mRNA-seq V2 Library Prep kit for Illumina. Paired-end sequencing was performed on the HiSeq X Ten system generating 2 x 150 bp reads.

Raw transcriptome reads were quality filtered using Trimmomatic version 0.33 (69) based on the following criteria: (1) remove any leading or lagging bases with quality scores less than 20, (2) clip off any Illumina adapters, (3) scan the read with a 5-base wide sliding window and cut when the average quality score per base drops below 20, and (4) remove any sequences shorter than 50 bp. The trimmed reads, both paired and unpaired, were then mapped to the WP2 genome using BBMap version 38.81 (69), with all default settings except for requiring a minimum identity for mapping of 90%. The mapped reads were then used to generate count tables of the number of transcripts that map to each gene using the featureCounts program version 1.6.3 (70) with default settings. Differential expression of genes was identified between different temperatures in each growth phase using DESeq2 version 1.22.2 (71). Genes were considered to be significantly differentially expressed if the log2 fold change between conditions was greater than 1 or less than -1 and if the adjusted p-value was less than or equal to 0.05. The gene expression data (**Fig. S1**) was visualized using the *geom_boxplot* function in the R package *ggplot2* (version 3.3.5) to present the median ratio normalization (MRN) values from replicate samples at each growth phase. If multiple genes were associated with a reaction, the expression levels of each gene were included as a data point in the box plot.

### Genome-scale metabolic reconstruction

A genome-scale model was constructed for WP2 based on ortholog mapping and manual curation. Protein sequences were downloaded from the GenBank Assembly Database for the full genomes of *S. psychrophila* WP2 (GCA_002005305.1), *S. piezotolerans* WP3 (GCA_000014885.1), *S. oneidensis* MR-1 (GCA_000146165.2), *Shewanella* sp. MR4 (GCA_000014685.1), *Shewanella* sp. W3-18-1 (GCA_000015185.1), and *S. denitrificans* OS217 (GCA_000013765.1). Orthologous genes between the genomes were identified using a bi-directional best hit BLASTp approach as detailed in a previous study (72). The *psammotate* function in the PSAMM software package version 1.1 (73) was used to generate a draft model of WP2 based on the predicted orthologous genes and previously published GEMs of *S. piezotolerans* WP3 (34), *S. oneidensis* MR-1 (68), *Shewanella* sp. MR4, *Shewanella* sp. W3-18-1, and *S. denitrificans* OS217 (74) (**Data Set S1, Tab 2**). Following the initial reconstruction, eggNOG-mapper (version 1) (75, 76) was used to assign putative functions to the WP2 genes based on the eggNOG database version 5.0 (77) and to map WP2 genes to the Kyoto Encyclopedia of Genes and Genomes (KEGG) (78) and COG (79) databases. These functional assignments were manually curated to identify functions and genes missing from the draft model. A biomass objective function for WP2 was constructed based on the genome sequence, predicted coding sequences, and predicted proteome of WP2 (37) (**Data Set S1, Tab 3**). The fatty acid composition of the WP2 model was calculated based on the experimentally measured lipid composition in WP2 (33), and the trace element component of the biomass was formulated based on the values used in the iJO1366 *E. coli* GEM (80). This biomass formulation was used for simulations performed at all three temperatures.

Metabolic gaps in the WP2 model were identified using the *fastgapfill* (*81*) and *gapfill* (*82*) functions as implemented in PSAMM version 1.1. The gap filling process was performed using the LMO-812 medium (34) with experimentally verified combinations of carbon sources and electron acceptors (33). Reactions from template models of other *Shewanella* species, the iJO1366 *E. coli* model (80), and the BIGG database version 1.5 (83) were used as references for the gap filling. Metabolic reactions predicted by the gap filling step were included in the model when a corresponding gene was identified in the WP2 genome. Gap reactions were added to the model without gene associations only if they were required for the model to produce biomass (**Data Set S1, Tab 4**). Charge and mass balance in the WP2 model was verified using the *chargecheck*, *formulacheck*, and *masscheck* functions in PSAMM version 1.1 (73). Model stoichiometric consistency, mass balance, charge balance, and metabolite connectivity were further verified using the MEMOTE software (49). The complete WP2 GEM was compared to previously reported growth phenotypes for WP2 to confirm that it can utilize the experimentally verified carbon sources and electron acceptors for growth (**Data Set S1, Tab 1**).

### Identification of thermodynamic constraints

Standard Gibbs free energy change of reaction (*Δ_r_G’*^∘^) was calculated for reactions in the WP2 model using the Group Contribution python package (84) (**Data Set S1, Tab 5**). Multiple settings for the Group Contribution prediction were tested to evaluate the effects of the pH and ionic strength parameters on the predicted *Δ_r_G’*^∘^ values. Only small differences were observed in the predictions when different settings were used, and due to these properties being unknown in WP2, the standard condition for the *Δ_r_G’*^∘^ estimation was defined as pH 7, ionic strength of 0 M, and 25°C. In order to run the Group Contribution approach on individual reactions in the WP2 model, all metabolites participating in a reaction were represented in one of two forms: (1) identifiers in the Thermodynamics of Enzyme-Catalyzed Reactions Database (TECRdb) (85), which provides thermodynamic data for a collection of enzyme-catalyzed reactions; (2) metabolite structure represented as Simplified Molecular-Input Line-Entry System (SMILES) strings (**Data Set S1, Tab 6**), which can be used to derive information related to metabolite properties for the *Δ_r_G’*^∘^calculation. Metabolites in the WP2 model were first mapped to the TECRdb based on their names and formulas, and when a mapping was not available, metabolite structural information was collected from the Pubchem Database (86) in the form of the canonical SMILES strings. When SMILES strings were not available in the Pubchem database, for example, for chitin, eicosapentaenoic acid (EPA), 3-hydroxy-11-methyldodecanoic acid, 11-octadecenoic acid, and the fatty-acyl ACP compounds, metabolites structures were manually drawn using the MarvinSketch application version 19.11 (ChemAxon, https://www.chemaxon.com) and exported using the SMILES export function. Artificial metabolites representing macromolecule biomass components like DNA, RNA, and the overall biomass compound were not assigned SMILES strings. The *Δ_r_G*^∘^ of the biomass objective function, macromolecular biosynthesis reactions, and several reactions in the cell wall biosynthesis pathway were not predicted due to the complexity of the metabolites involved in these reactions.

### Configuration of the temperature-dependent models

The TMFA approach (44, 45) was applied to the WP2 model using the *tmfa* function implemented in PSAMM (version 1.1) in python version 3.7.10. All simulations were done with the global maximum allowable flux being set to 100 and the minimum allowable flux being set to -100. The PSAMM *tmfa* implementation takes as input a configuration file that references the *Δ_r_G’*^∘^estimations as described in a prior section (**Data Set S1, Tab 5**), a list of reactions excluded from the *Δ_r_G’*^∘^calculation due to the high complexity of the involved metabolite’s structures (**Data Set S1, Tab 7**), a list of transport reactions mapped to parameters (*i.e.*, net charge and net protons transported from outside to inside of the cell) that can be used to calculate energy associated with electrochemical potential and pH gradient across the cell membrane (**Data Set S1, Tab 8**), and definitions of metabolite concentrations and exchange constraints (**Data Set S1, Tab 9 and 10**). The temperature-dependent simulation of the WP2 model was achieved by introducing a number of constraints that are specific to the growth profiles of the organism under suboptimal, optimal, and supraoptimal temperatures. These included the calculation of *Δ_r_G’*values in the TMFA simulation using the temperature in Kelvin, the identification of temperature-dependent ATP maintenance constraints, and the constraint of oxygen concentrations and exchange fluxes based on the solubility of oxygen at each given temperature.

By default, the concentrations of all metabolites were constrained to be within a range of 1e-5 to 0.02 mol/L following prior studies (44, 87), with the exception of metabolites present in the experimental media and a pair of metabolites involved in an essential metabolic reaction (**Data Set S1, Tab 9**). First, the concentrations of two metabolites, n-carbamoyl-L-aspartate and (S)-dihydroorotate, were expanded with a range from 1e-6 to 0.05 mol/L following prior conventions [38], to enable thermodynamic feasibility in an essential dihydroorotase (DHORTS) reaction. Second, the upper bound of oxygen concentration was set according to the solubility of oxygen in liquid media under different temperatures. Next, the compounds that represent nutrient inputs in the LMO-812 media had the upper bound of their concentrations assigned based on the composition of the media. Finally, the lower bound of media components that had a concentration lower than 1e-4 mol/L was set to two orders of magnitude lower than its upper bound to allow for some variability in the concentrations. The pH in the model was constrained between 6 and 8 for both the cytosolic and extracellular compartments. The exchange flux constraints for the TMFA simulation were formulated to represent the availability of nutrients and the diffusion of metabolic products (**Data Set S1, Tab 10**). The exchange constraints of carbon, nitrogen, sulfur, and phosphorus sources were constrained based on experimental media. Small molecules and metal ions were released to enable free exchange, and the lower bound of oxygen exchange was set according to the solubility of oxygen in liquid media under different temperatures.

An ATPM reaction was added to each model to account for the non-growth associated ATP maintenance (**Fig. 1**). The ATPM reaction was individually calibrated for each temperature-dependent model based on the experimentally measured growth rates. First, a robustness analysis was performed by successively increasing the flux of ATPM while determining the optimized biomass over each step. Next, a linear fitting was performed using the ATPM flux and the biomass flux values. Lastly, the upper and lower bounds of the ATPM reaction were determined by projecting the experimentally measured growth rates based on the linear fitting (**Fig. 1B-D**). A range was determined for the ATPM constraint for each temperature-dependent model based on the range of growth rates observed in the experimental replicates of each temperature.

### Random simulations with temperature-dependent, thermodynamically constrained models

Metabolic simulations were performed on each temperature-dependent Model with the same growth media as in the WP2 growth experiments, using 5 mM of GlcNac as the sole carbon source. To probe for the utilization of metabolic pathways under different temperatures, metabolic reaction fluxes, Gibbs free energy change of reactions (*Δ_r_G’*), and metabolite concentrations were predicted by running each model through 1,000 random simulations. Each simulation was performed by constraining the biomass flux to a random value selected within the range of experimentally determined growth rates ([0.110, 0.112] for 4°C; [0.254, 0.255] for 15°C; and [0.042, 0.135] for 20°C). The resulting predictions of reaction fluxes, *Δ_r_G’*, and metabolite concentrations were summarized by calculating the minimum, maximum, median, and 25th and 75th percentiles over the 1,000 random simulations for each temperature-dependent model (**Data Set S2**). Significant changes among the temperature-dependent models were identified across the 1,000 random simulations for each metric based on the Kruskal-Wallis test and corresponding pairwise post hoc tests, using the *kruskal.test* and the *pairwise.wilcox.test* function, respectively, in the R package *stats* (version 4.1.3). The effect size was calculated using the *epsilonSquared* function in the R package *rcompanion* (version 2.4.15). Comparisons of the 4°C, 15°C, and 20°C models with effect size □ 0.36, following previously established thresholds (88), and with p-value □ 0.05 in both Kruskal-Wallis test and pairwise tests were reported as carrying significant changes across the different temperatures (**Data Set S2**).

The CUE was calculated from each random simulation using the equation 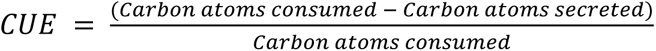 following an existing approach (50), where all the carbon-containing exchange compounds were grouped into lists of consumed or produced metabolites based on the direction of their exchange fluxes in a simulation. The exchange fluxes were scaled based on the number of carbon atoms in each compound, and sums of the scaled fluxes were taken for the list of consumed and produced metabolites, respectively, to obtain the *carbon atoms consumed* and the *carbon atoms secreted* parameters in the CUE equation.

To calculate the total amount of ATP produced or consumed in a model simulation, the metabolic flux of each reaction that involved ATP was multiplied by the stoichiometry of ATP in that reaction. The resulting positive values indicated the production of ATP, while negative values indicated the consumption of ATP. The flux of the ATPM reaction was excluded from these calculations so that the ATP production and consumption values reported in this study reflected the energy requirements related specifically to the metabolic processes. The ATP production per unit of carbon source was calculated through dividing the sum of all stoichiometrically-scaled ATP-producing fluxes by the exchange flux of the carbon source (*i.e.*, GlcNac). The ATP consumed per gDW biomass was calculated through dividing the sum of all stoichiometrically-scaled ATP-consuming fluxes by the biomass flux.

The metabolite ratios (**Fig. 4**) were calculated based on the metabolite concentrations predicted in each random simulation. The visualization of metabolite ratios (**Fig. 4**), metabolic efficiency metrics (**Fig. 2**), and individual metabolic fluxes (**Fig. S1**) was created with the *geom_boxplot* function in the R package *ggplot2* (version 3.3.5), using collections of values obtained from the 1,000 random simulations. Pairwise comparison of *Δ_r_G’*, metabolic flux, and gene expression values across different temperature conditions (**Fig. S2**) were visualized with scatter plots using the *geom_point* function in *ggplot2* (version 3.3.5), with linear fittings visualized with the *stat_smooth* function using the method *lm*. The *Δ_r_G’*and metabolic flux were scaled for each reaction across three temperature conditions. This was done by dividing each metric by the root mean square of the same metric across three temperature conditions. The expression values were normalized by MRN and scaled for each individual gene by the root mean square values across three temperatures.

## Data and software availability

Modeling approaches implemented in this study are accessible through the open-source PSAMM software, release v1.1 and are freely available in a git repository: https://github.com/zhanglab/psamm. The WP2 GEM in both YAML and SBML formats, all inputs used for the modeling, and all analysis scripts are available on GitHub at the following address: https://github.com/zhanglab/GEM-iWP2. Transcriptome raw read data was deposited in the NCBI Sequence Read Archive database under bioproject PRJNA739408 (https://dataview.ncbi.nlm.nih.gov/object/PRJNA739408?reviewer=d79hr02aj87t57hbotv8mnc27i).

## Supplemental Figure Legends

**Fig. S1. Temperature-dependent metabolic fluxes and corresponding gene expressions in the key steps of WP2 central metabolism.** Metabolic fluxes predicted from the 4°C (blue), 15°C (yellow), or 20°C (red) models are shown as box plots based on values obtained from 1,000 random simulations. Significant differences between fluxes were identified using the Kruskal-Wallis test (**Materials and Methods**) and are marked by brackets with an asterisk. Gene expression data are shown separately for the early exponential phase (early), the late exponential phase (late), and the stationary phase (stat). Counts obtained from median ratio normalization (MRN) were presented in a log scale for the comparison of gene expression levels across different temperatures. The expression values are marked as significantly different if any of the genes associated with the reactions showed greater than 1 or less than -1 log2 fold change in expression and had an adjusted p-value of less than 0.05 based on the differential expression analysis (**Materials and Methods**).

**Fig. S2. Comparisons of** *Δ_r_G’***, metabolic flux, and gene expression under different temperature conditions.** The *Δ_r_G’* (top row) and metabolic flux (middle row) values were obtained from simulations of the temperature-dependent models, scaled for each reaction across the three temperature conditions (**Materials and Methods**). The gene expression (bottom row) was based on MRN values, scaled for each gene across the three temperature conditions (**Materials and Methods**). The scatter plots demonstrate pairwise comparisons of the scaled values between each pair: 4°C versus 15°C (left), 4°C versus 20°C (center), 15°C versus 20°C (right). The linear fitting was shown as a blue line in each plot, with gray shading showing the 95% confidence interval. The corresponding linear equations and r^2^ values were shown with black labels.

## Supplemental Data Legends

**Data Set S1: Metabolic model information and constraints used for model simulations. Tab 1)** Comparison of model-predicted and experimentally measured growth and no-growth Phenotypes. G indicates that growth was observed (experimental) or predicted (model). NG indicates that no growth was observed (experimental) or predicted (model). **Tab 2)** Ortholog mapping between WP2 and four previously modeled *Shewanella* species. Reactions associated with each gene are provided for each of the previously modeled strains. **Tab 3)** The composition and stoichiometry of each biomass component and the overall biomass reaction in the WP2 GEM. **Tab 4)** Gap reactions included in WP2 GEM. The reason for including the gap reaction and source are listed for each reaction. **Tab 5)** Standard Gibbs Free Energy predictions for reactions in the WP2 GEM based on the Group Contribution method. Values are provided in kJ/mol. **Tab 6)** Metabolite structural information. Table contains metabolite names, formulas, and charges from the model. SMILES strings were provided along with an indication of the source of the structure. A mapping to the TECRdb database used by the Group Contribution package was also included. **Tab 7)** List of all reactions that were excluded from the thermodynamic constraints. **Tab 8)** Net number of protons and net charge transported from outside of the cell to inside of the cell for each transport reaction in the WP2 GEM. **Tab 9)** Non-default concentration bounds used in the WP2 TMFA simulations. Concentrations were based on the LMO-812 media composition for all compounds except cpd_cbasp and cpd_dhor-S which were modified based on previous literature (Materials and Methods). **Tab 10)** Exchange Constraints used for the temperature-dependent model simulations based on the LMO-812 media.

**Data Set S2. Summary of TMFA simulation results. Tab 1)** Summarized values for reaction fluxes from random TMFA simulations. The median, minimum, maximum, 25th percentile, and 75th percentile values were included for each reaction based on the 1,000 random TMFA simulations. Results of a Kruskal-Wallis test, estimate of the effect size, and results of pairwise Wilcoxon tests are shown for each reaction. Values of ND are reported when the 3 distributions being compared have no difference. **Tab 2)** Summarized values for reaction Gibbs free energy values from random TMFA simulations. The median, minimum, maximum, 25th percentile, and 75th percentile values were included for each reaction based on the 1,000 random TMFA simulations. Results of a Kruskal-Wallis test, estimate of the effect size, and results of pairwise Wilcoxon tests are shown for each reaction. Values of ND are reported when the 3 distributions being compared have no difference. **Tab 3)** Summarized values for metabolite concentrations (shown as the natural log of the concentration) from random TMFA simulations. The median, minimum, maximum, 25th percentile, and 75th percentile values were included for each reaction based on the 1,000 random TMFA simulations. Results of a Kruskal-Wallis test, estimate of the effect size, and results of pairwise Wilcoxon tests are shown for each metabolite. Values of ND are reported when the 3 distributions being compared have no difference.

**Data Set S3. Summary of differential gene expression analyses. Tab 1)** Differential expression of genes between temperatures during early exponential growth phase. **Tab 2)** Differential expression of genes between temperatures for late exponential growth phase. **Tab 3)** Differential expression of genes between temperatures for stationary growth phase.

